# Performance-optimized hierarchical models only partially predict neural responses during perceptual decision making

**DOI:** 10.1101/221630

**Authors:** Laura Gwilliams, Jean-Rémi King

## Abstract

Models of perceptual decision making have historically been designed to maximally explain behaviour and brain activity independently of their ability to actually perform tasks. More recently, performance-optimized models have been shown to correlate with brain responses to images and thus present a complementary approach to understand perceptual processes. In the present study, we compare how these approaches comparatively account for the spatio-temporal organization of neural responses elicited by ambiguous visual stimuli. Forty-six healthy human subjects performed perceptual decisions on briefly flashed stimuli constructed from ambiguous characters. The stimuli were designed to have 7 orthogonal properties, ranging from low-sensory levels (e.g. spatial location of the stimulus) to conceptual (whether stimulus is a letter or a digit) and task levels (i.e. required hand movement). Magneto-encephalography source and decoding analyses revealed that these 7 levels of representations are sequentially encoded by the cortical hierarchy, and actively maintained until the subject responds. This hierarchy appeared poorly correlated to normative, drift-diffusion, and 5-layer convolutional neural networks (CNN) optimized to accurately categorize alpha-numeric characters, but partially matched the sequence of activations of 3/6 state-of-the-art CNNs trained for natural image labeling (VGG-16, VGG-19, MobileNet). Additionally, we identify several systematic discrepancies between these CNNs and brain activity, revealing the importance of single-trial learning and recurrent processing. Overall, our results strengthen the notion that performance-optimized algorithms can converge towards the computational solution implemented by the human visual system, and open possible avenues to improve artificial perceptual decision making.

## 1 Introduction

Perception - deriving an orderly categorization of the outside world - is a foundational problem being addressed by both artificial intelligence (AI) and cognitive neuroscience. In the last decade, AI systems have proved capable of resolving perceptual labeling at a level competitive to biological systems. However, whether AI and biological algorithms use the same computational strategies remains an open question.

Factors that makes categorization so challenging are invariance (one physical object can lead to multiple distinct sensory inputs) and ambiguity (identical sensory inputs can be consistent with more than one physical object). For example, any two-dimensional projection of the world on the retina is by definition compatible with an infinite number of three-dimensional objects. Yet, despite of these challenges, humans solve perceptual inference seemingly effortlessly.

What are the computational principles upholding this ability? Three lines of research, each echoing Marr’s tri-level epistemology, have historically tackled this issue. At the computational level, a vast body of literature converge on the notion that simple perceptual decisions, including illusory ones, follow Bayesian inference principles, whereby reports match the most plausible causes of ambiguous sensory evidence [15]. At the implementational level, electrophysiology and neuroimaging converge on the idea that perceptual recognition, should it be of simple visual objects [11], alphabetic characters [9] or faces [5] depend on dedicated hierarchies of neural assemblies that incrementally combine increasingly abstract features.

However, conclusions concerning Marr’s intermediary - “algorithmic”- are less agreed upon. On the one hand, system neuroscience has generated a rich set of simple models to account for perceptual decision making. For example, Shadlen and collaborators have shown how the pyramidal neurons in the lateral intra-parietal cortex mimic drift-to-bound models [22] to integrate lower-level evidence over time [6]. Complementarily, Dehaene and collaborators have argued that the fronto-parietal cortices implement a winner-take-all attractor to select the representation that will ultimately guide behavior [3]. These “late-resolution” models can be contrasted to computational architectures that resolves noise and ambiguity with recurrent processes at the early visual stages [17] and/or throughout distributed feedforward hierarchies [15, 23]. On the other hand, AI research has led to a great variety of computational architectures, based on normative [16], shallow, deep and recurrent architectures [18], each with competitive efficiency.

Each of these computational architectures can be thought of as a candidate algorithm to explain how the brain performs perceptual judgments. Algorithms can be distinguished in terms of the representations they compute at each processing stage, and can thus be compared to the contents of neuronal activity at each instant.

In the present study, we aimed to identify, from MEG recordings, the computational architecture supporting perceptual decision making in human subjects presented with variably ambiguous characters. To this aim, we designed minimalistic stimuli that varied along 7 orthogonalized dimensions. Each dimension specifically targeted one of the transformations putatively required by our perceptual decision making paradigm. We then decoded each of these stimulus properties from brain activity, and compared their time courses to those of multiple shallow and deep architectures tested on the same task.

## 2 Method

### Procedure

Three experiments, consisting of identifying briefly flashed ambiguous characters, were performed by three distinct groups of subjects. In experiments 1 and 2, 12 and 17 subjects performed a subjective or an implicit categorization task, respectively. 10 euros was provided as compensation, and each study took 1 hour. In experiment 3, 17 subjects performed an identification task inside an Elekta Neuromag MEG scanner and were given 70 euros as compensation. All experiments were approved by the local ethics committee. All subjects signed an informed consent form.

### Stimuli

Stimuli were displayed on a 50% gray background at (60 Hz refresh rate) as controlled with Psychtoolbox [21]. The overall visibility of the stimuli was artificially diminished via backward masking (Experiment 1), overall contrast manipulation (Experiment 2) or crowding (Experiment 3) to enhance decision making processes. Ten non-ambiguous stimuli (characters 0, 4, 5, 6, 8, 9, H, E, C, A) were used as a base across all experiments, and were composed of 4 to 7 identical edges horizontally or vertically oriented. Ambiguous stimuli were constructed from a linear combination of any two characters distinguished by a single edge (e.g. 4-H). Eight levels of morph-contrast, linearly distributed between 0 and 1 were used.

### Tasks

Experiments 1 and 2 served to ensure that the characters were indeed being perceived categorically. In experiment 1, subjects were invited to provide a continuous report; responses were categorically distributed around the non-ambiguous stimuli, suggesting a categorical percept (Figure 1, left). In experiment 2 subjects indicated whether two simultaneously flashed stimuli were identical or not. We hypothesize that if stimuli are automatically perceived categorically, then two highly ambiguous stimuli will be more often judged as different than two weakly ambiguous stimuli. Our results confirmed this hypothesis (Figure 1, middle).

**Figure 1:**
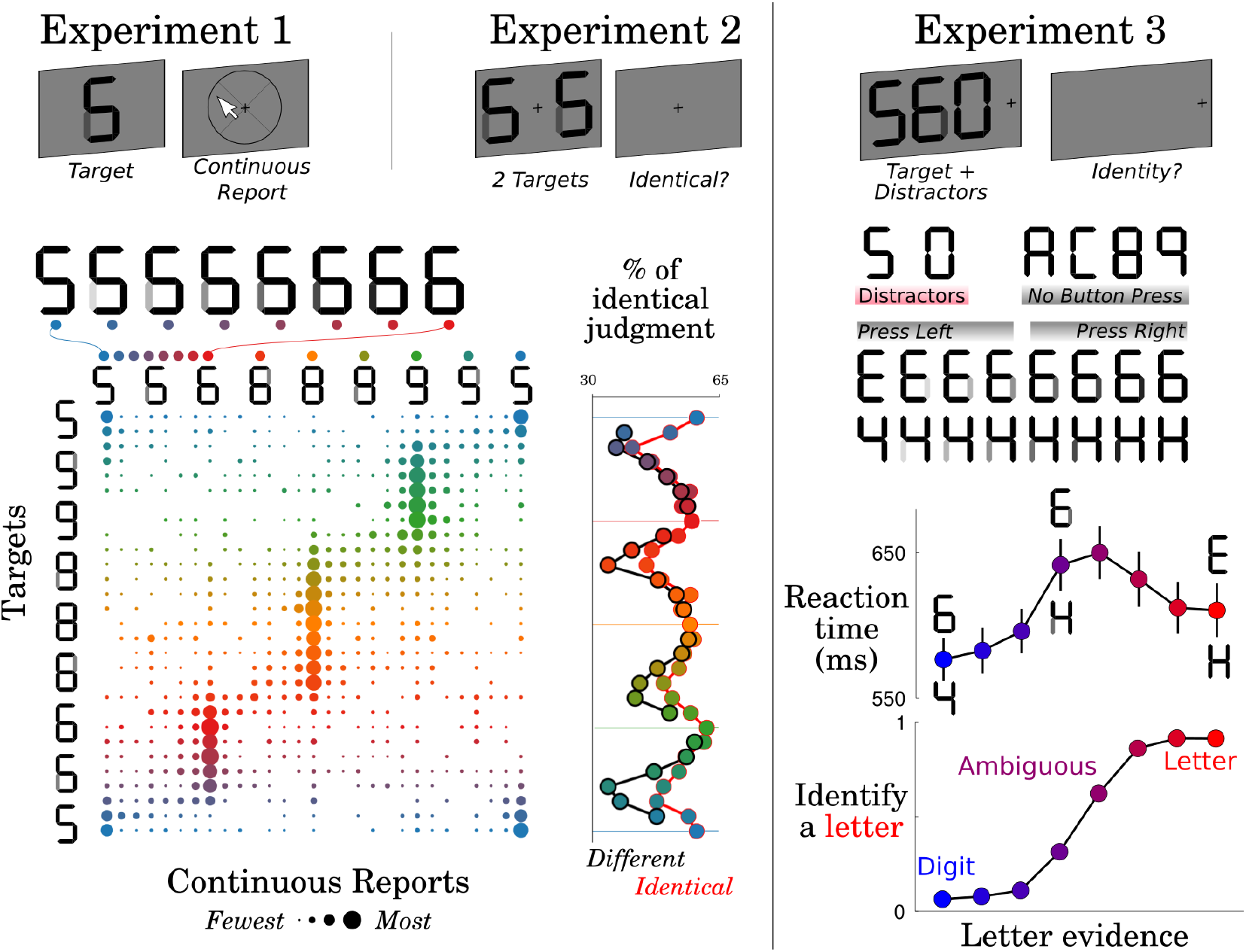
Experiment 1: A variably ambiguous target stimulus was flashed on the screen and judged with continuous perceptual reports. The matrix depicts the proportion of trials (disk size) rated with each perceptual judgement (x-axis) for each of the 28 possible ambiguous stimulus (y-axis). The results show that subjects tended to report ambiguous stimuli as their closest categorical stereotype. Experiment 2: A pair of identical or different stimuli was rapidly flashed and had to be judged as “identical” or not. The proportion of “identical” ratings diminished with the stimuli ambiguity, suggesting automatic categorical perception. Experiment 3: A crowded target, variably morphed between a letter (red) and a digit (blue), was rapidly flashed onto the screen and had to be identified with a block-specific identity-response mapping. Subjects correctly identified targets as a function of the objective evidence, and slow down reaction times for ambiguous targets.

In Experiment 3, brain activity was recorded with MEG while subjects were presented with letters (e.g. E, H, A, C), digits (6, 4, 8, 9), or mixture between letters and digits. The target was surrounded by two constant distractors, and presented as a brief flash lasting 100 ms either on the left or on the right of fixation (Figure 1, right). On 83% of the trials, subjects were instructed to press, within 2 seconds, a button with their left hand or right hand to indicate the identity of the stimulus. This identity-response mapping changed at every block to orthogonalize the putative conceptual category (‘letter’, ‘digit’) and from the motor response (‘left’, ‘right’). On the other 17% of the trials, subjects only had to pay attention to the stimuli, they did not have to indicate their percept. A total of 1920 trials, grouped into 40 blocks, were performed by each subject.

### Neuroimaging

Anatomical MRIs (3T Siemens MRI scanner with 1x1x1.1 mm voxels) were acquired after the MEG acquisition. Grey and white matter were segmented with Freesurfer and coregistered with each subject’s digitized head shapes along with three fiducial points. The raw MEG data was corrected with Maxfilter’s SSS, bandpass filtered between 0.5 and 40 Hz (MNE default parameters with firwin design). Epochs were cut from −300 ms to +1500 ms time-locked to stimulus onset, and −1000 to +500 time-locked to the motor response and finally downsampled to 250 Hz. The forward solution for source localisation was computed separately for each subject, which maps source space to sensor space given the brain anatomy of the subject. An inverse operator was computed from the forward solution and the noise covariance over sensors averaged over all trials. The inverse model was then applied to each single-trial epoch, assuming an SNR of 3, loose dipole fitting at 0.2, normal orientation of the dipole relative to the cortical sheet. The result was a dynamic Statistical Parameter Map (dSPM) [2] value at each vertex of the reconstruction, for each millisecond of the epoch. Decoding analyses were performed within a 5-split stratified cross-validation and using 12-regularized *z*-scored linear estimators fit across all MEG sensors recorded at a unique time sample. Continuous variables were fit with a ridge regression (alpha=1), scored with a Spearman R. Categorical variables were fit with a logistic regression (C=1) and scored with the area under the curve (AUC). Temporal generalization was performed by testing whether the estimator fit at a given time sample could accurately predict other time samples [14], respectively aiming to detect the effects of (i) stimulus position, (ii) the total number of edges presented on the screen, (iii) the contrast of the critical feature, (iv) the continuous evidence in favor of a letter, (v) whether the stimulus should be categorized as a digit or a letter, (vi) the difficulty of the decision and (vi) the actual response button pressed by the subjects. By design, all but (iv) and (v) were orthogonalized (see Figure 4A), and could thus be decoded independently of one another. Statistical effects are based on second-level spatio-temporal cluster-testing with 10,000 permutations) across subjects. All analyses were performed using MNE [7], scikit-learn [20] and Scipy [12] with default parameters.

### Models

Four types of computational models were trained and/or tested to perform the task of experiment 3. Model 1 is single-layer model, input with 14 features, corresponding to each of the visual edges manipulated in experiment 3 and consists of identifying the Bayesian-optimal combination of feature that maximally predicts each of the task-features. Model 2 is a 5-layer convolutional neural network (2 convolutional layers (32, (3x3) relu), followed by one max pooling (2x2) and 25% dropout, and ending with two successive dense (128 relu, 36 softmax) layers separated by a 50% dropout, optimized with ADAM on the categorical cross-entropy across 36 classes (26 letters and 10 digits) using 72,869 images generated a single character generated from 1,000 fonts presented on the left or right side of the image and tested on the ambiguous stimuli presented to the subjects. The third type of model consists of VGG-16, VGG-19, ResNet-50, Xception, InceptionV3, and MobileNet. For concision purposes, we presently focus on VGG-19, which we will refer to as Model 3 [25], trained on ImageNet to label 1,000 classes from 1M images, and ResNet-50, which we will refer to as Model 4 [10], also trained on ImageNet with the same labelling task. Note these models are not optimized to perform our task, and were tested with the very same images presented to subjects.

**Figure 2:**
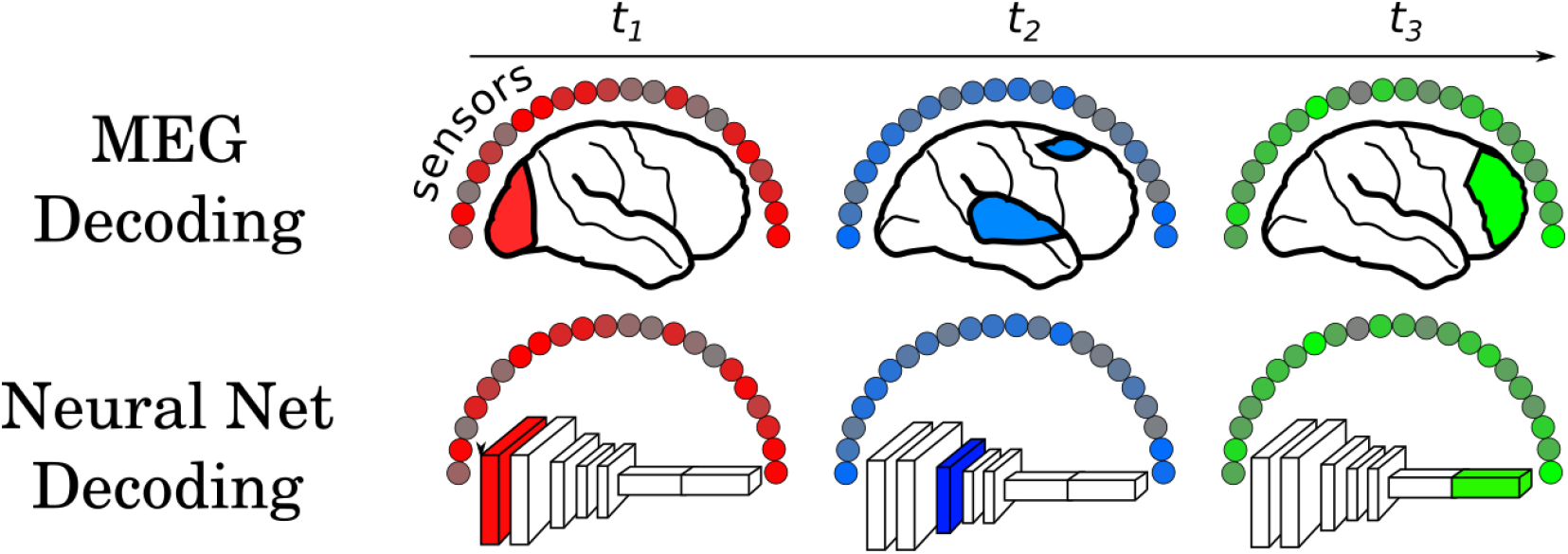
Analogous decoding analyses were applied to the MEG and neural network data. Top: A classifier is trained independently at each millisecond of the brain data across all 306 MEG sensors. Bottom: A classifier is trained at each layer of a neural network, from artificial neurons that have been projected into a 306-dimensional space using random Gaussian projections. In both cases, the classifiers were fit to decode each of 7 properties of the stimuli. Temporal generalization was applied to test how well a classifier fit at one time-sample / computational layer was able to predict activity at other time-points / layers. This can be used to estimate the maintenance of each feature representation.

## 3 Results

### 3.1 The brain progressively transforms ambiguous characters into categorical representations

Seventeen subjects had to identify a rapidly flashed variably-ambiguous character onto the left or the right side of fixation, while their brain activity was recorded with MEG (Figure 1). The task was designed to make the response buttons orthogonal to 6 other task-related features, ranging from low-level sensory features (stimulus position, number of edges composing the stimulus) to perceptual (contrast of the ambiguous edge) and conceptual level (evidence and optimal decision in favor of a letter character). Subjects’ behavior appeared typical of decision-making tasks: the psychometric function followed a sharp sigmoidal pattern and reaction time increased in ambiguous (645 ms) as opposed to non-ambiguous items (580 ms).

We implemented source and decoding analyses at each time sample following the stimulus onset to determine whether and when the brain activity was different between letters and digits. Low, but strongly significant letter/digit decoding could be observed from ~150 ms up until 940 ms with a peak around 370 ms (AUC=55%, p < .001). These results confirm that human subjects automatically encode characters as abstract letter and digit categories even when the task does not require them to. Source reconstruction suggest that these neural codes were generated in the ventral and dorsal stream similarly to previous electro-corticography and fMRI research [24, 9], but were too variable across subjects to survive correction for multiple comparison.

We then investigated whether and how the brain transforms ambiguous visual inputs into abstract letter/digit categories. Specifically, we hypothesized that just a simple remapping of the stimulus would be characterized by a linear correlation between the brain activity and objective evidence in favor of the letter category. By contrast, a categorical representation would be non-linearly correlated with the evidence: All stimuli that are more letter-like that digit like would be treated the same, and the brain would not be sensitive to within-category variance. Using multiple regression on the probabilistic predictions of the letter/digit classifiers, our results show that the neural responses linearly correlate with the evidence between ~200 and 400 ms after stimulus onset, and subsequently correlate with the optimal categories. Single-layer, recurrent and deep architectures designed to mimic these neural responses suggest that only recurrent and deep neural networks can progressively transition from a linear to a categorical mapping of ambiguous stimulus (Figure 3). Temporal generalization (TG) analyses can be used to distinguish our recurrent and deep architecture, by predicting square and diagonal TG matrices respectively. TG analyses of the MEG data appeared diagonal across all decodable time samples, thus suggesting that the categorization of ambiguous stimuli is supported by a deep feedforward neural architecture. Overall these results suggest that the disambiguation algorithm of the human brain is performed by a long series of processing stages typical of deep neural networks.

**Figure 3:**
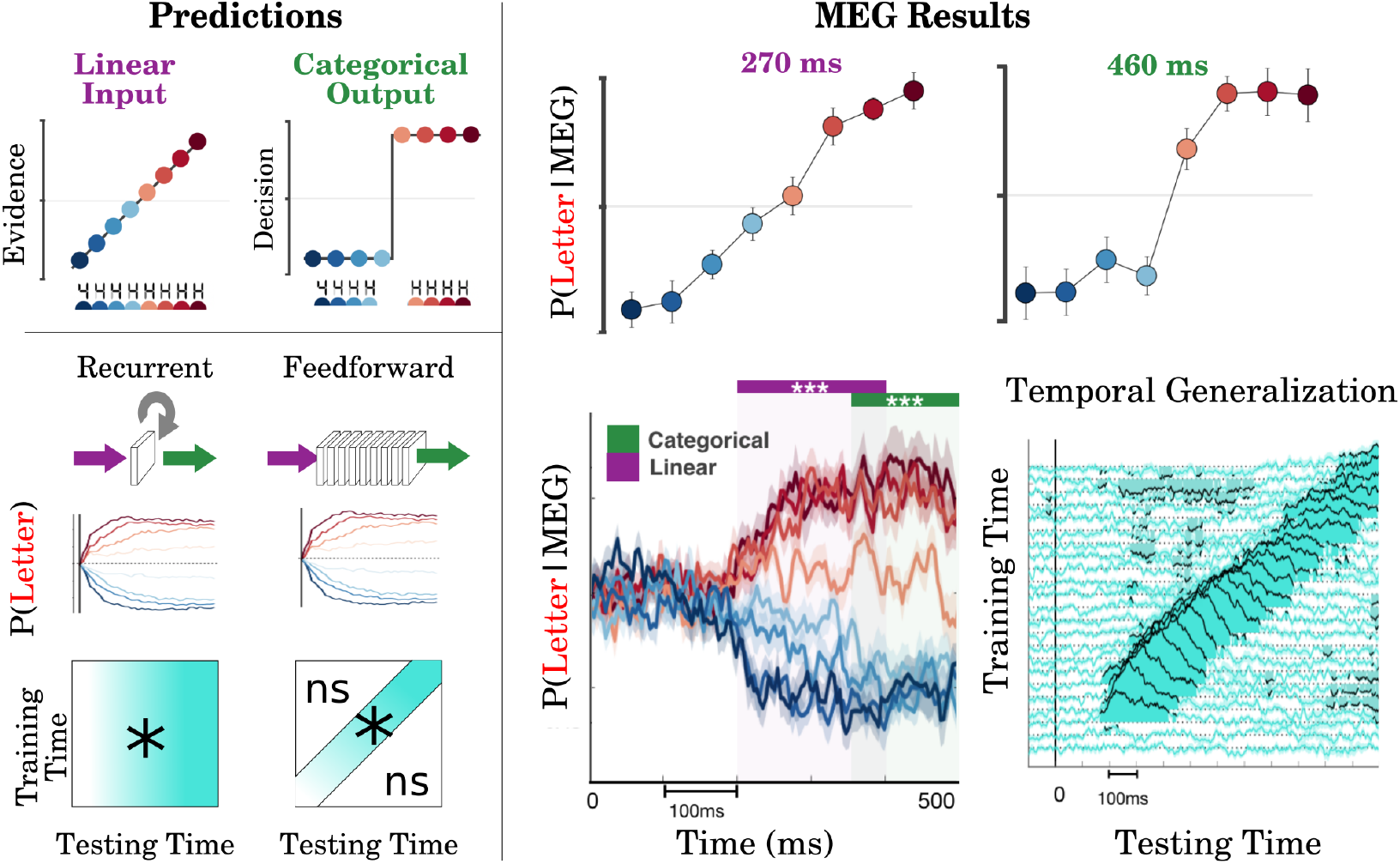
Left: Simple perceptual decisions can be modeled as a non-linear transformation (green) of a variably-ambiguous (purple) stimulus. Both recurrent and deep neural network architectures can progressively transform an ambiguous input into a category, leading to drift-diffusion like responses over time: i.e. activations that ramp towards the most likely category at a speed proportional to the evidence for that category. Simple recurrent networks are characterized by a stable spatio-temporal profile leading to a square temporal generalization (TG) matrix. Simple feedforward architectures can be associated with a constant change of activations and thus lead to a diagonal TG matrix. Right: Probabilistic estimates of letter versus digits estimated from MEG signals at each time sample show that the neural activity correlates linearly with the evidence from 200 to 400 ms, and then becomes categorical (400-500 ms). TG analyses of the MEG (turquoise) reveal a diagonal patterns typical of deep neural networks.

### 3.2 The brain sequentially transforms sensory inputs to task-related representations

To test whether perceptual decision making was comprised of a series of processing stages in the human brain, we implemented source and decoding analyses at each time sample following the stimulus onset and isolated 7 putative levels of representations characterizing each stimulus. The results show that all of the seven features can be decoded from the MEG evoked responses (Figure 4). Critically, each representation started to be decodable and peaked sequentially as a function of their putative level of abstraction: brain activity was first modulated by stimulus position (mean peak: 120 ms, AUC=94%), then the number of edges (peak: 250 ms; R=16%), contrast (peak: 340 ms, R=7%), Letter/Digit evidence and category (peak: 370 ms, R=12%, AUC= 55%), and ended with the difficulty (peak: 590 ms, R=8%) and response button (peak: 600 ms, AUC=65%). All of these features remained decodable up until subjects’ response. Overall, these results suggest that the human brain hierarchically encodes the sensory stimulus into a series of increasingly task-oriented representations, reminiscent of cascade models [19]. The temporal generalization analyses (Figure 4D) suggest that most of the stimulus properties are processed in a feedforward manner, as indicated by the diagonal generalization pattern. However, not all features could be accounted for with a purely feedforward architecture: The TG matrix for stimulus side shows reactivation patterns, supporting the presence of recurrent processing for this feature.

**Figure 4:**
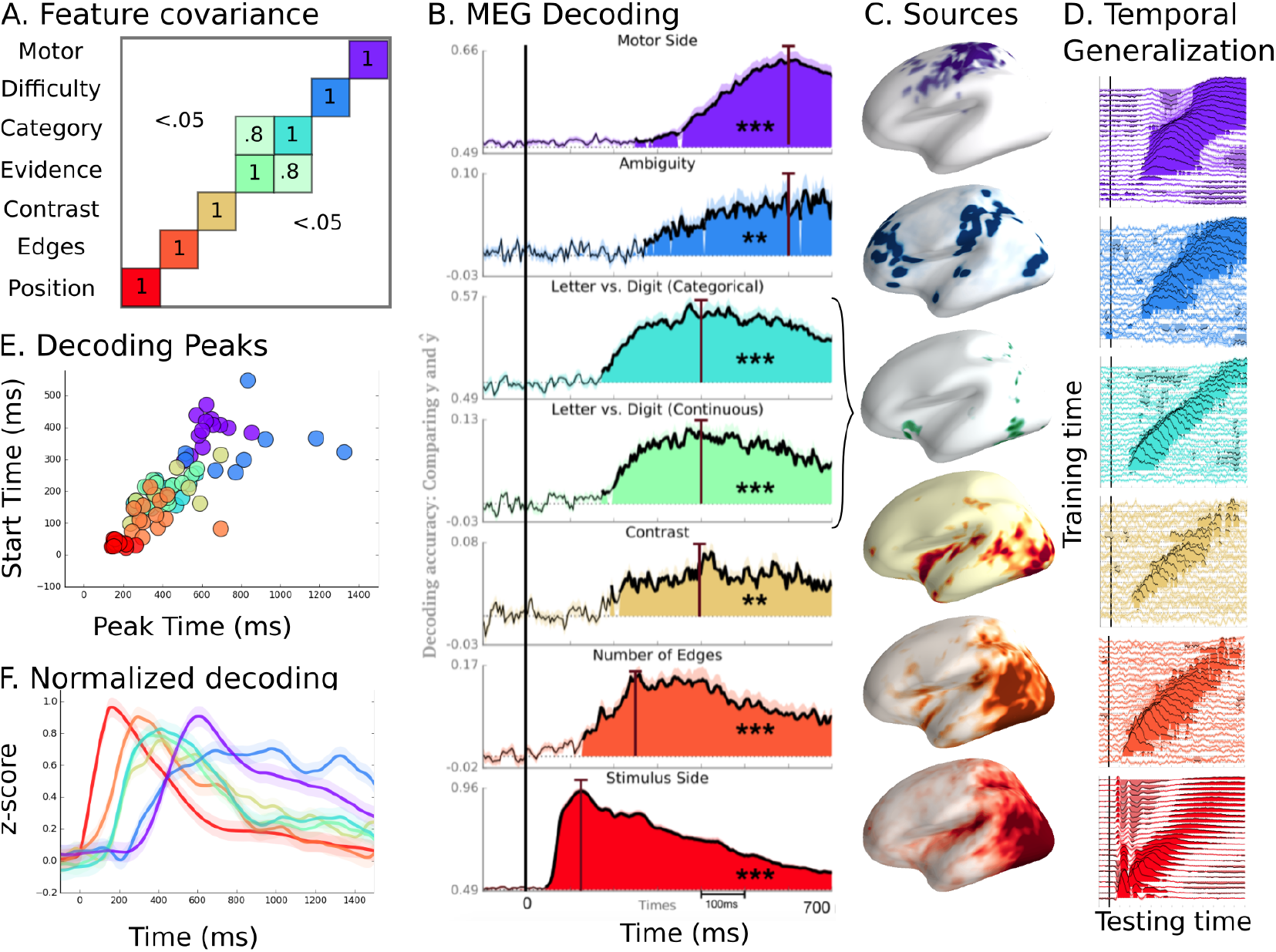
A: The covariance matrix indicates how each of the seven properties covary with one another (R^2^). B: Decoding scores (y-axis) for each of the seven features, when MEG classifiers are trained and tested at the same milliseconds (x-axis) respective to stimulus onset. Colored areas indicate statistical significance across subjects. The vertical lines indicate the time of each decoding peak. C: MEG source localization at peak time. D: Temporal generalization analysis consists of training a classifier at each time-point, and testing it at all other time-points, in order to assess the maintenance of each representation. Diagonal patterns suggest that the coding of neuronal activity changes over time. E: Time at which decoding starts (y-axis) and peaks (x-axis) for individual subjects. F: Normalized decoding scores. Overall, the results demonstrate that each of the features is sequentially encoded in the brain activity, in a way that specifically matches efficiency-optimized CNNs.

### 3.3 Distinct computational architectures can equally perform character categorization, but predict different latent representations

The computational models were designed as descriptions of neural responses, and not trained to efficiently solve the task. To better probe the computational architecture underlying perceptual decision making, we thus compared brain responses to four feedforward computational models that all performed our perceptual decision task at (or close to) optimality. The models differed in their overall architecture as well in the task for which they were optimized. Importantly they all predict distinct latent representations decodable at each processing stage.

Our results demonstrate that a single softmax layer, taking the 14 visual edges as inputs can already accurately categorize all of the target stimuli of experiment 3 (Model 1). Multivariate decoding of its inputs efficiently extracts all but the ‘difficulty’ task features. In other words, given the right features, a single layer model can optimally perform our task, but predict that all but the difficulty feature should be equally and simultaneously decodable.

We then trained a series of shallow and deep convolutional neural networks (i) trained either on character recognition (Model 2) or image labeling (Models 3 & 4), and (ii) tested with our task by replacing the final softmax layer fit for our task. Since the input features are pixels, and thus oversample the inputs of Models 1, models 2, 3 and 4 subsume single-layer model. To identify the specific predictions of these deeper models, we thus randomly projected the hidden activations of each layer onto 306 virtual Gaussian sensors. This operation, repeated 17 times, aimed at mimicking the within-subject decoding approach undertaken with MEG, and constrains the to-be-decoded features to elicit an activation in a sufficiently high proportion of neurons to be picked by a small number of random projections. Our results showed that the optimized CNN (Model 2) adequately identify all characters (>94% accuracy, chance=1/36th), and adequately label the stimuli designed for our task (100% accuracy). Additionally, the results suggest that we can equally decode the seven orthogonalized features of our task at all layers of the neural network.

Finally, we input our stimuli to the CNNs trained for natural image labeling (Models 3 & 4), and analyzed them similarly to Model 2. Although these neural networks (VGG-19 and ResNet-50) were not trained for character recognition, we could decode of each of the task-features. However, and contrary to the other models, each of the 7 features could not be equally decoded from all layers. First, early layers could only be used to decode the position of the stimulus. Second, the decoding of the number of edges progressively increased with depth. Third, the perceptual (contrast) conceptual (letter evidence and category) and motor response could only be decoded from the deep, fully connected layers. Finally, once a feature could be decoded in a given layer, it remained equally decodable within each of the subsequent layers. One of the other models we tested (MobileNet) also showed this sequential decoding pattern, whereas the remaining models we tested, Xception and Inception V3, did not.

This pattern of results is neither compatible with single layer models (Models 1), nor with our shallow CNN (Model 2) optimized for character recognition. By contrast, this sequence of neural responses strongly correlate with the hierarchy of latent representation generated by deep neural networks trained for complex natural-image labeling, at the notable exception of the difficulty feature.

**Figure 5:**
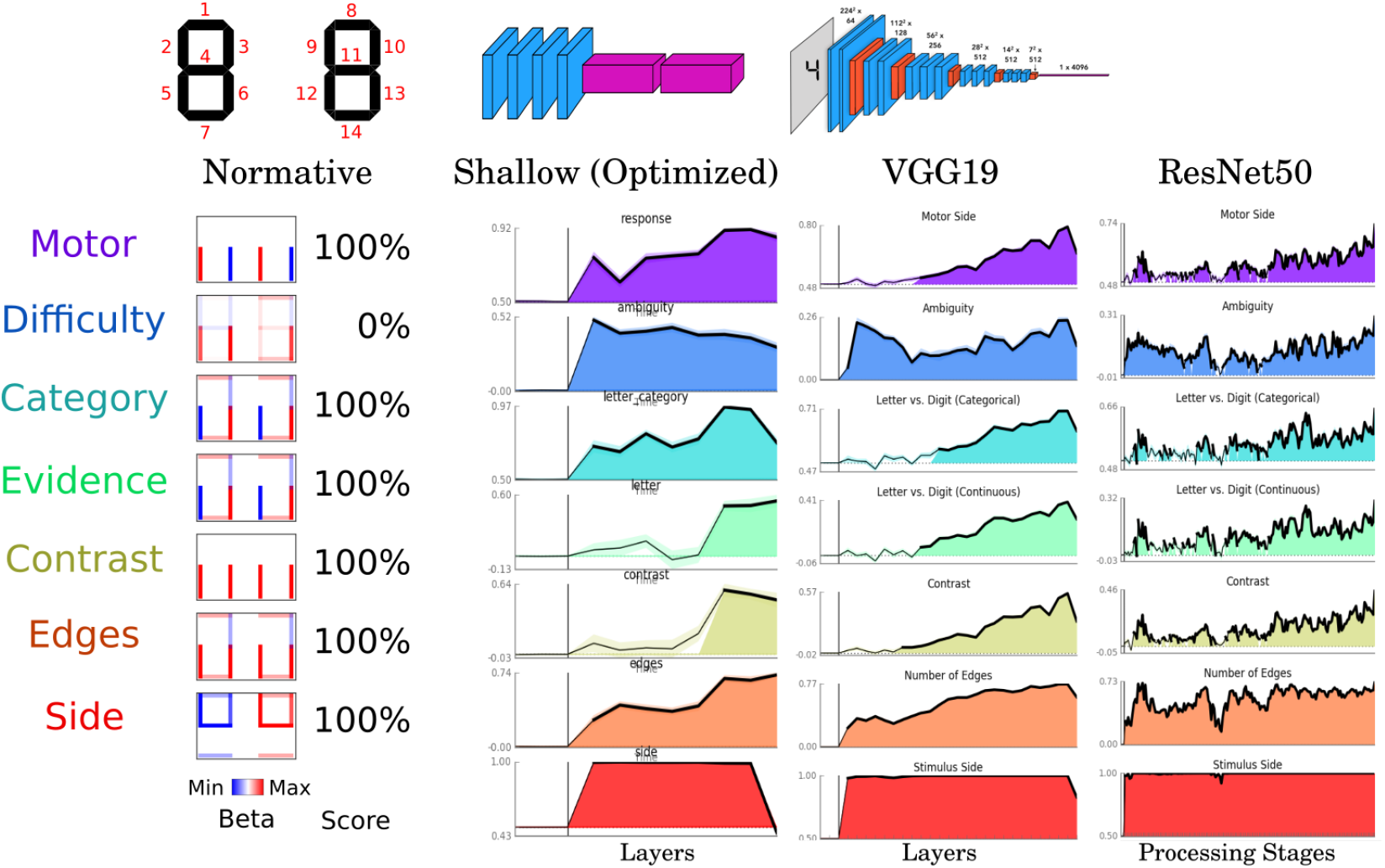
Decoding performance for each of the tested computational models. The normative model distinguishes stimulus features based on optimal linear combinations of the 14 visual edges that make up the stimulus input. All features were perfectly decodable, other than the input difficulty. The shallow CNN was trained to identify a character label (e.g. “H”, “E”, “4”) from a wide range of letter and digit characters. It was not explicitly taught the distinction between the letter/digit category. The decoding plot is analogous to Figure 4 for the MEG data in terms of the available features at each processing stage. VGG19 is a 19-layer CNN optimized to label thousands of categories from natural images. ResNet50 is a 50-layer CNN trained similarly to VGG19. Each model was able to perform the task, but predicted different hidden representations between the image and the decision.

## 4 Discussion

Both behavioral and MEG analyses confirmed that subjects automatically categorize ambiguous characters, seemingly thanks to weak but detectable neural assemblies of the ventral and dorsal cortical hierarchies as previously reported by fMRI [9] and electro-corticography [24]. Importantly, the representations estimated from letter/digit probabilistic classifiers, revealed typical drift-diffusion patterns [22] and thus echoes the classic findings in the electrophysiology of perceptual decision making [6]. Indeed, our results appear similar to the classic ramping activity observed by Shadlen and collaborators in the LIP of macaque monkeys during perceptual decision making tasks: the single trial predictions were decodable from ~200 ms, ramped at a speed proportional to the objective evidence manipulated in a given trial, and up until a subject’s motor action.

However, source and temporal generalization analyses suggest that the transformation of ambiguous evidence into a categorical representation is not encoded in the activity of a unique neural assemblies. Rather, ambiguous representations are sequentially and repeatedly re-coded via a long series of transient and increasingly categorical neural assemblies, a particular computational feature also found in deep as well as in some recurrent architectures. The possibility that human perceptual decision making depends on such architecture is further strengthened by the specific correlation observed between (i) the decoding time courses of the 7 orthogonalized features estimated from the brain activity and (ii) those estimated from the deep neural networks.

However, two lines of results suggest that none of the presently tested models fully account for brain activity. First, the “difficulty” feature systematically appeared decodable from the early processing stages of our models; by contrast, in the MEG data, this feature only rises late (simultaneous to the motor responses) and lasted until the beginning of the subsequent trials. This suggests that the “difficulty” feature may be more relevant to learning than sensory extraction. Indeed, unlike the presently-tested models, the human brain is known to adjust its decision parameters at each trial [26]. The second source of discrepancy comes from source and temporal generalization analyses, which, contrary to our feedforward models, suggest that some neural codes are partially maintained over time. This particular result suggests that recurrent and/or top-down connections need to be added to our computational models to better match the sequence of brain responses.

Overall, the present work strengthens a recent line of electrophysiology and neuroimaging research demonstrating several elements of convergence between the mammalian visual system and deep convolutional neural networks trained for image categorization [27, 8, 1, 4, 13]. Additionally, our results pave the way to better identify the computational architecture of human perception, and apply it to current advances in AI research.

## Acknowledgments

This project received funding from the European Union’s Horizon 2020 research and innovation program under grant agreement No 660086, the Bettencourt-Schueller Foundation, the Fondation Roger de Spoelberch, the Philippe Foundation and the Abu Dhabi Institute G1001. We are infinitely grateful to Stanislas Dehaene for his support. We would also like to thank Christelle Larzabal, Denis Engemann, and Sabrina Medalel for their help.

## References

[1] R. M. Cichy, A. Khosla, D. Pantazis, A. Torralba, and A. Oliva. Comparison of deep neural networks to spatio-temporal cortical dynamics of human visual object recognition reveals hierarchical correspondence. Scientific reports, 6:27755, 2016.

[2] A. M. Dale, A. K. Liu, B. R. Fischl, R. L. Buckner, J. W. Belliveau, J. D. Lewine, and E. Halgren. Dynamic statistical parametric mapping: combining fmri and meg for high-resolution imaging of cortical activity. Neuron, 26(1):55–67, 2000.

[3] S. Dehaene.and J.-P. Changeux. Experimental and theoretical approaches to conscious processing. Neuron, 70(2):200–227, 2011.

[4] M. Eickenberg, A. Gramfort, G. Varoquaux, and B. Thirion. Seeing it all: Convolutional network layers map the function of the human visual system. NeuroImage, 152:184–194, 2017.

[5] W. A. Freiwald and D. Y. Tsao. Functional compartmentalization and viewpoint generalization within the macaque face-processing system. Science, 330(6005):845–851, 2010.

[6] J. I. Gold and M. N. Shadlen. The neural basis of decision making. Annu. Rev. Neurosci., 30:535–574, 2007.

[7] A. Gramfort, M. Luessi, E. Larson, D. A. Engemann, D. Strohmeier, C. Brodbeck, L. Parkkonen, and M. S. Hämäläinen. Mne software for processing meg and eeg data. Neuroimage, 86:446–460, 2014.

[8] U. Güçlü and M. A. van Gerven. Deep neural networks reveal a gradient in the complexity of neural representations across the ventral stream. Journal of Neuroscience, 35(27):10005–10014, 2015.

[9] T. Hannagan, A. Amedi, L. Cohen, G. Dehaene-Lambertz, and S. Dehaene. Origins of the specialization for letters and numbers in ventral occipitotemporal cortex. Trends in cognitive sciences, 19(7):374–382, 2015.

[10] K. He, X. Zhang, S. Ren, and J. Sun. Deep residual learning for image recognition. In Proceedings of the IEEE conference on computer vision and pattern recognition, pages 770-778, 2016.

[11] D. H. Hubel and T. N. Wiesel. Receptive fields, binocular interaction and functional architecture in the cat’s visual cortex. The Journal of physiology, 160(1):106–154, 1962.

[12] E. Jones, T. Oliphant, and P. Peterson. {SciPy}: open source scientific tools for {Python}. 2014.

[13] S.-M. Khaligh-Razavi and N. Kriegeskorte. Deep supervised, but not unsupervised, models may explain it cortical representation. PLoS computational biology, 10(11):e1003915, 2014.

[14] J. King and S. Dehaene. Characterizing the dynamics of mental representations: the temporal generalization method. Trends in cognitive sciences, 18(4):203–210, 2014.

[15] D. C. Knill and A. Pouget. The bayesian brain: the role of uncertainty in neural coding and computation. TRENDS in Neurosciences, 27(12):712–719, 2004.

[16] B. M. Lake, T. D. Ullman, J. B. Tenenbaum, and S. J. Gershman. Building machines that learn and think like people. Behavioral and Brain Sciences, pages 1–101, 2016.

[17] V. A. Lamme and P. R. Roelfsema. The distinct modes of vision offered by feedforward and recurrent processing. Trends in neurosciences, 23(11):571–579, 2000.

[18] Y. LeCun, Y. Bengio, and G. Hinton. Deep learning. Nature, 521(7553):436–444, 2015.

[19] J. L. McClelland. On the time relations of mental processes: An examination of systems of processes in cascade. Psychological review, 86(4):287, 1979.

[20] F. Pedregosa, G. Varoquaux, A. Gramfort, V. Michel, B. Thirion, O. Grisel, M. Blondel, P. Prettenhofer, R. Weiss, V. Dubourg, et al. Scikit-learn: Machine learning in python. Journal of Machine Learning Research, 12(Oct):2825–2830, 2011.

[21] D. G. Pelli. The videotoolbox software for visual psychophysics: Transforming numbers into movies. Spatial vision, 10(4):437–442, 1997.

[22] R. Ratcliff and J. N. Rouder. Modeling response times for two-choice decisions. Psychological Science, 9(5):347–356, 1998.

[23] M. Riesenhuber and T. Poggio. Hierarchical models of object recognition in cortex. Nature neuroscience, 2(11), 1999.

[24] J. Shum, D. Hermes, B. L. Foster, M. Dastjerdi, V. Rangarajan, J. Winawer, K. J. Miller, and J. Parvizi. A brain area for visual numerals. Journal of Neuroscience, 33(16):6709–6715, 2013.

[25] K. Simonyan and A. Zisserman. Very deep convolutional networks for large-scale image recognition. arXiv preprint arXiv:1409.1556, 2014.

[26] C. Summerfield. Dissociable sources of uncertainty in perceptual decision making. PhD thesis, University of Oxford, 2016.

[27] D. L. Yamins, H. Hong, C. F. Cadieu, E. A. Solomon, D. Seibert, and J. J. DiCarlo. Performance-optimized hierarchical models predict neural responses in higher visual cortex. Proceedings of the National Academy of Sciences, 111(23):8619–8624, 2014.

